# Spermidine dampens inflammation by directly inhibiting Th17 cytokine production through a PRDX1 associated antioxidant pathway

**DOI:** 10.1101/2021.08.31.458409

**Authors:** Jiadi Luo, Yong Joon Kim, Xiaojing An, Li Fan, Carla Erb, Dequan Lou, Yong Yao, Annabel A. Ferguson, Yinghong Pan, Kong Chen

## Abstract

The activation of IL-17 signaling has been linked to the pathogenesis of many chronic, inflammatory lung diseases including Cystic Fibrosis (CF). Through unbiased single-cell RNAseq screening, we found that IL-17+ T cells highly express *Srm* and *Smox*, which encode two key enzymes for spermidine synthesis. Spermidine has been shown to reduce inflammation by regulating macrophage activation and balancing Th17/Treg differentiation, but its direct effects on Th17 cytokine production has not been carefully investigated. Here, using already differentiated Th17 cells from cultured mouse splenocytes, we found that exogenous spermidine directly inhibits IL-1β/IL-23 induced IL-17 production. Blockade of endogenous spermidine synthesis enhanced IL-17 production above native levels, further supporting that spermidine is a direct regulator of cytokine secretion independent of differentiation. *In vivo*, spermidine alleviates lung inflammation in both PA infection and LPS induced acute lung injury models. Further RNA-seq analysis suggests spermidine suppression of Th17 cytokine production is mediated through its PRDX1 dependent antioxidant activity. Our data establishes that spermidine is a direct regulator of Type-17 T cell cytokine production and has potent anti-inflammatory effects against lung inflammation.

## Introduction

The Th17/IL-17 signaling pathway is an important regulator of immune and inflammatory function. Th17 cells play a key role in host immunity, particularly in combating infection and maintaining mucosal tissue barrier function.^1, 2, 3, 4, 5^ During activation of the Th17/IL-17 pathway, Th17 cells produce IL-17 family cytokines that bind specifically to downstream IL-17 receptors, leading to the release of other inflammatory cell chemokines and stimulating further inflammatory cascades. The Th17/IL-17 pathway thus plays an important immune activating role. Pathologically excessive activation of the Th17/IL-17 response, however, has been implicated in autoimmune and inflammatory disorders such as psoriasis, ankylosing spondylitis, multiple sclerosis, and systemic inflammatory response syndrome (SIRS).^1^

Although Th17/IL-17 signaling cascade has been extensively implicated in various physiologic and pathologic contexts, the regulatory mechanisms of the Th17/IL-17 pathway remain unclear. Further delineation of the regulatory mechanism of the Th17/IL-17 signaling pathway is needed to guide clinical management of IL-17 mediated inflammation and autoimmunity.

In this study, we unexpectedly found that spermidine synthase (SRM) and spermine oxidase (SMOX) are highly expressed in *Il17a*+ cells. SRM and SMOX are the most important catalytic enzymes for the synthesis of spermidine (Spd) from putrescine and spermine, respectively.^6, 7^ Spermidine is a polyaliphatic amine with anti-inflammatory properties.^8,9,10,11,12^ Recent reports highlighted that spermidine plays important roles in regulating T-cell lineage differentiation.^13,14,15^ However, the direct impact of spermidine on Th17 cytokine production has not been carefully examined. In addition, Spd has been linked to a variety of anti-inflammatory mechanisms but has not been directly linked to the regulatory pathway for Th17/IL-17 cytokine secretion prior to this study.

Here we establish that Spd directly regulates the Th17/IL-17 signaling pathway in already differentiated T cells including memory T cells and gamma-delta T cells. The regulatory roles of spermidine and its inhibitors in the Th17/IL-17 pathway were characterized *in vitro* and *in vivo* using RNA sequencing, RT-PCR, ELISA, and other technical means. We found that Spd can directly target T cells and downregulate the expression of Th17/IL-17 signaling pathway products, thereby inhibiting IL-17 cytokine-induced inflammation. Conversely, inhibition of Spd synthesis can reverse Spd’s suppressive effects on inflammatory cytokine release in the Th17/IL-17 pathway. Further single-cell RNA-seq (scRNA-Seq) results revealed that Spd’s regulation of cytokine production depends on the antioxidant activity of peroxiredoxin 1 (PRDX1) in Th17 cells. The results of this study establish Spd’s regulatory role in Th17/IL-17 signaling, providing further guidance for the potential pharmacologic management of inflammation and hyper-immunity.

## Results

### Enzymes for spermidine synthesis (Srm, Smox) are upregulated and Spermidine is downregulated upon Th17/IL-17 pathway activation in Th17 cells

To determine the molecular pathways that may be implicated in Th17/IL-17 signaling, we carefully mined a dataset where mouse peritoneal lavage cells were stimulated with IL-1β/IL-23 and assessed with scRNA-Seq, with a focus on 4 subsets of T-cell only (Figure 1A). Surprisingly, *Srm* and *Smox* genes were found to be upregulated in IL-17+ expressing cells (Figure 1A). Spermidine synthase (Srm) and spermine oxidase (Smox) are both catalytic enzymes for spermidine synthesis. To further explore the relationship between spermidine and Th17/IL-17 pathway, we assessed *Srm* and *Smox* expression levels with qPCR in spleen cells and splenic T cells derived from adult mice. Adult mice were used for their matured immune systems with already differentiated Th17 cells, allowing for the investigation of spermidine’s direct role on T cells independent of the differentiation regulatory pathway.

**Figure 1.**
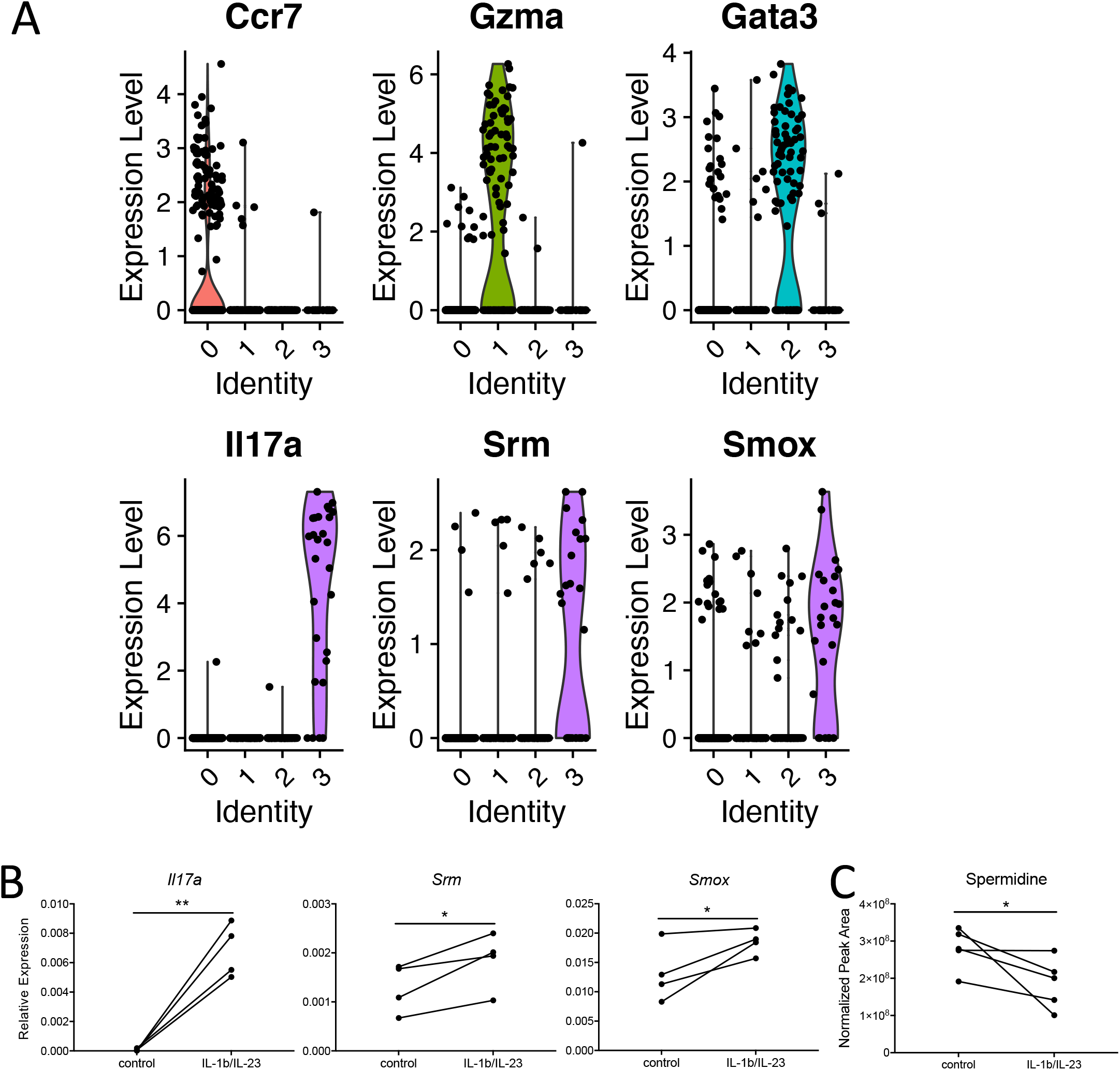
Polyamine metabolism is involved in IL-17 production. A: Mouse peritoneal lavage cells induced by thioglycollate were collected and stimulated with IL-1b/IL-23, and scRNA data confirmed IL-17 producing cells have high *Srm* and *Smox* expression. B and C: Fresh enriched spleen T cells were stimulated with mouse IL-1b +IL-23 50ng/ml overnight, (B) *Il17a*, *Srm* and *Smox* were highly induced, detected by qPCR, gene expression relative to Gapdh (graphs displayed in panel B are representative ones from 4 repeated experiments, N=2-4 in each experiment); (C) Polyamine in the cytoplasm of T cells were measured by HPLC system.

RT-PCR analysis confirmed that *Srm* and *Smox* are highly expressed in IL-17 producing cells upon stimulation by IL-1β/IL-23 (Figure 1B).

Next, to further explore the link between spermidine and Th17 cytokine production, we directly measured the levels of spermidine and other polyamines (spermine and putrescine) in T cells after Th17/IL-17 pathway activation. Despite the upregulation of spermidine synthesis enzymes, lower spermidine levels were observed after Th17/IL-17 pathway stimulation (Figure 1C) whereas other polyamines remained unchanged (results not shown). The *Srm* and *Smox* gene expression profile and changes in spermidine levels in mature T cells suggest that spermidine may be involved in the regulation of the Th17/IL-17 pathway independent of its regulatory role in T cell differentiation.

### Spermidine directly targets T cells and down-regulates Th17/IL-17 pathway products *in vitro*

To determine the direct effect of spermidine on the Th17/IL-17 pathway, the cytokine expression profile of activated T cells was measured in the presence of exogenous spermidine. We monitored cytokine regulation by measuring mRNA expression (qPCR) and protein levels (ELISA) for Th17 effector cytokines.

Upon administration of spermidine, both the gene expression (Figure 2A) and protein levels (Figure 2B) of Th17/IL-17 cytokine products were downregulated in T cells stimulated with IL-1β/IL-23. We also observed downregulated gene expression and cytokine levels for the Th1 effector cytokine, IFN-γ, which is typically activated in conjunction with the Th17/IL-17 pathway when fighting foreign infections.^16,17^ These results indicate that spermidine acts directly on T cells in downregulating Th17/IL-17 pathway products. Upon administration of Difluoromethylornithine (DFMO), an inhibitor of spermidine synthesis^18^, the expression of Th17/IL-17 pathway product genes *Il17a*, *Il17f*, *Il22* (Figure 2C) and IL-17A protein (Figure 2D) were enhanced above IL-1β/IL-23 stimulation alone. DFMO did not, however, affect the IFN-γ pathway (Figure 2C & 2D). This result reinforces that spermidine may be one of the key native regulatory points specific to the Th17/IL-17 pathway.

**Figure 2:**
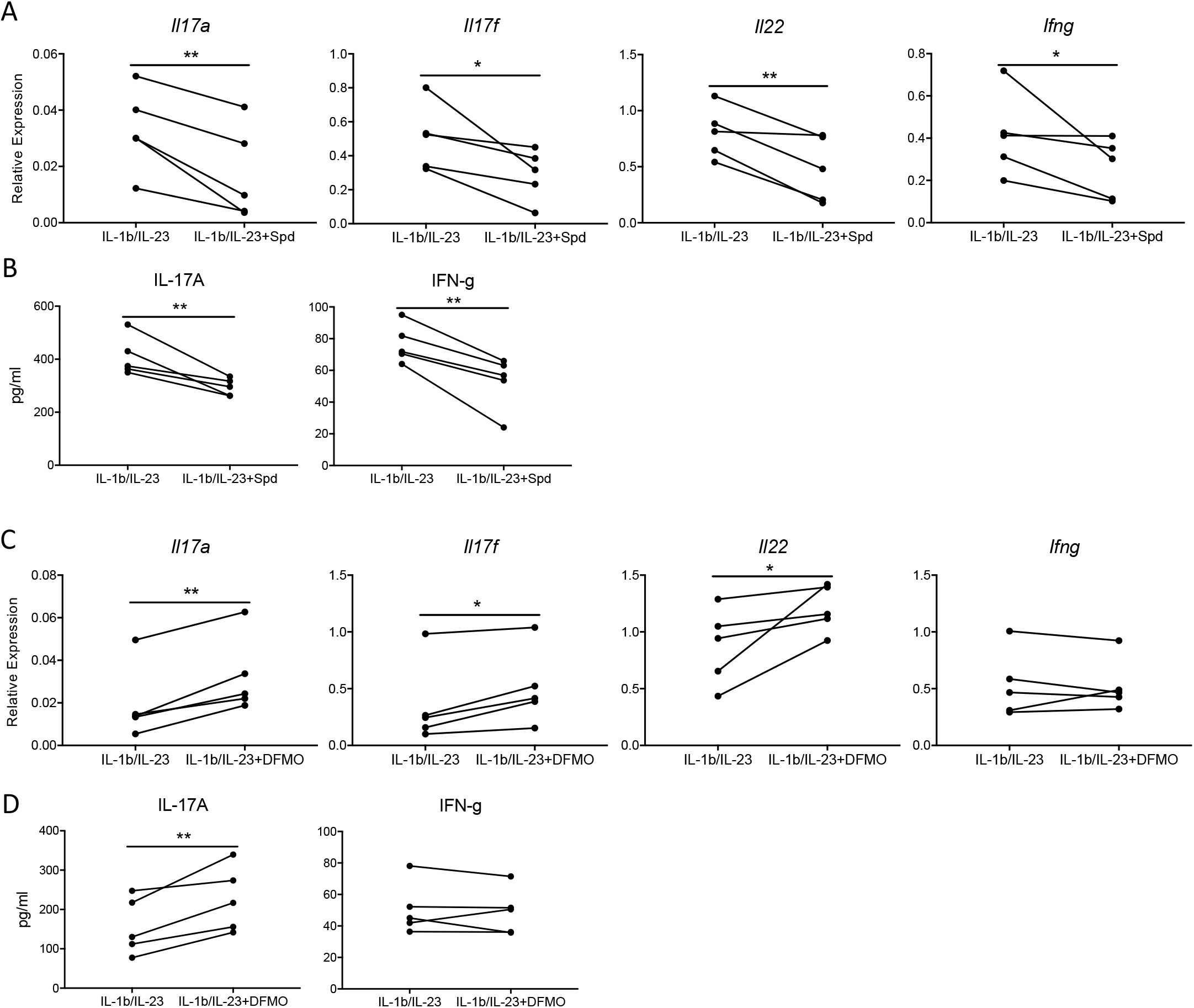
Spermidine downregulates the Th17/IL-17 by directly targeting T cells in vitro. Fresh enriched spleen T cells were stimulated with mouse IL-1b +IL-23 50ng/ml at the presence or absence of 3uM Spermidine or DFMO treatment overnight. This is a representative graph of five experiments, N=2-4 in each experiment. A: *Il17a*, *Il17f*, *Il22* and *Ifng* gene expression was measured by qPCR B: Relative expression to *Hprt*; IL-17A and IFN-g proteins were tested by ELISA. C: *Il17a*, *Il17f*, *Il22* and *Ifng* gene expression was measured by qPCR relative to *Hprt* expression D: IL-17A and IFN-g proteins were tested by ELISA

### Spermidine reduces pneumonia progression in LPS induced acute lung injury mouse model by downregulating the Th17/IL-17 pathway

With spermidine’s role in the negative regulation of Th17/IL-17 signaling established *in vitro*, we assessed spermidine’s effects on Th17/IL-17 signaling using an acute lung injury (ALI) mouse model. The ALI model is based on the context of bacterial pneumonia, where B6 WT mice were inoculated with LPS to induce lung inflammation and injury, with and without coadministration of spermidine. In this acute injury mouse model, spermidine downregulated Th17/IL-17 pathway expression, including *Il17a*, *Il17f* and *Il22* genes (Figure 3A) and IL-17A cytokine levels (Figure 3B). At the same time, the IFN-γ pathway, which is closely associated with the Th17/IL-17 pathway as previously described, was also inhibited by spermidine with decreased expression of the *Ifng* gene (Figure 3AB). Inhibition of native spermidine synthesis with DFMO resulted in the upregulation of Th17/IL-17 pathway genes (*Il17a*, *Il17f*, and *Il22*) as well as IL-17A protein expression (Figure 3CD). Consistent with the corresponding *in vitro* results, DFMO did not affect IFN-γ levels *in vivo* (Figure 3C & 3D).

**Figure 3.**
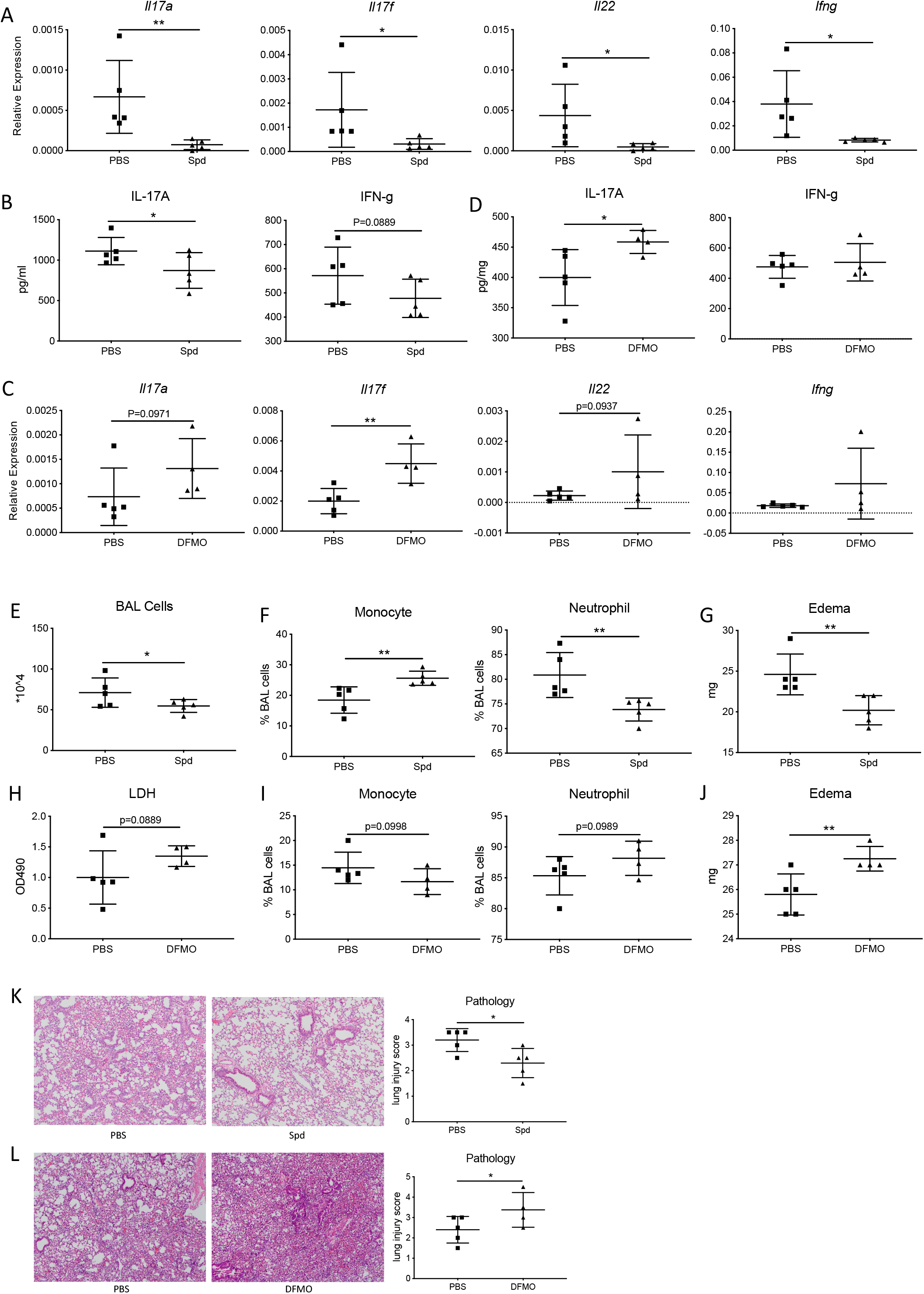
Spermidine dampens inflammation through inhibiting IL-17 production in an acute lung injury (ALI) *in vivo* model. N=5 in both PBS and Spd group. A-B: Lung tissue *Il17a* and *Ifng* mRNA (C) and protein (D) productions were obtained by ELISA and qPCR, relative expression to *Hprt*. C-D: Blockage of spermidine synthesis by DFMO in ALI mice model (N=5 in PBS group; N=4 in DFMO group). IL-17 family production at mRNA level (C) and protein level (D) in lung tissue were obtained by ELISA and qPCR, relative expression to *Hprt*. E: BAL cells were counted in PBS and spermidine treated lungs from ALI mouse models. F: BAL cell differentiation including monocyte and neutrophil were identified and quantified under the microscope after Kwik-Diff Staining. G: Right middle lung lobe was incubated in a 65C incubator for 48h, wet/dry weight ratio was recorded and edema levels were then calculated by wet weight minus dry weight H: DFMO administration resulted in higher levels of LDH, indicating more inflammation mediated cytotoxicity. I: DFMO group displayed less Monocytes and more Neutrophils in BAL cells. J: DFMO group displayed more severe lung edema, as calculated before in Figure 3G. K: Further lung tissue histology injury score was assessed. Inflammation can be seen in both groups. Compared to the PBS group, Spermidine group displayed less inflammatory cells, slighter alveolar space congestion, and less alveolar wall damage/alveolar fusion. This experiment has been repeated once. L: Further lung tissue histology injury score was assessed. Compared to the PBS group, DFMO group revealed more inflammatory cells infiltration, stronger alveolar space congestion and higher injury score. This experiment has been repeated once.

We assessed the degree of lung inflammation and damage in this mouse model and found that spermidine treated mice had less severe inflammation and lung damage (Figure 3E-G). There was an overall fewer number of immune cells in the BALF of the spermidine treated group (Figure 3E), indicating lower overall inflammation levels. Immune cells from spermidine administered lungs had higher proportions of nuclear cells and lower proportions of neutrophils (Figure 3F). In addition, edema levels were reduced in spermidine treated groups (Figure 3G). These measurements collectively indicate lower levels of inflammation in the lungs as a result of spermidine treatment. Not only did spermidine result in lower degrees of inflammation, but spermidine also protected against lung damage according to lung histology (Figure 3K). Inhibiting endogenous spermidine synthesis with DFMO promoted lung inflammation and injury, resulting in higher cytotoxicity, proportionally higher neutrophil counts, and elevated levels of lung edema (Figure 3H-J). Elevated levels of injury and inflammatory infiltration in DFMO treated lungs were also evident in lung histology (Figure 3L). Overall, the combination of lower expression of Th17/IL-17 pathway products and protected lung phenotypes suggest that spermidine protects against lung inflammation and injury by downregulating the Th17/IL-17 signaling pathway.

### Spermidine downregulates the Th17/IL-17 pathway in an acute live bacterial infection mouse model

A live bacterial infection mouse model was used to assess lung bacterial clearance upon spermidine modulation of the Th17/IL-17 pathway. It is well established that the body’s response to KP (*Klebsiella pneumoniae*) and PA (*Pseudomonas aeruginosa*), two gram-negative bacilli, are mainly mediated by the Th17/IL-17 pathway.^19,20^ In a PA infection model, the spermidine treatment group inhibited Th17/IL-17 pathway expression (Figure 4AB) but did not affect PA bacteria clearance (Figure 4C). Spermidine down-regulated Th17/IL-17 pathway expression, including *Il17a*, *Il17f* and *Il22* genes (Figure 4A) and IL-17A protein expression (Figure 4B). There was no difference in BALF protein concentration between the spermidine treatment group and the control group (results not shown).

**Figure 4:**
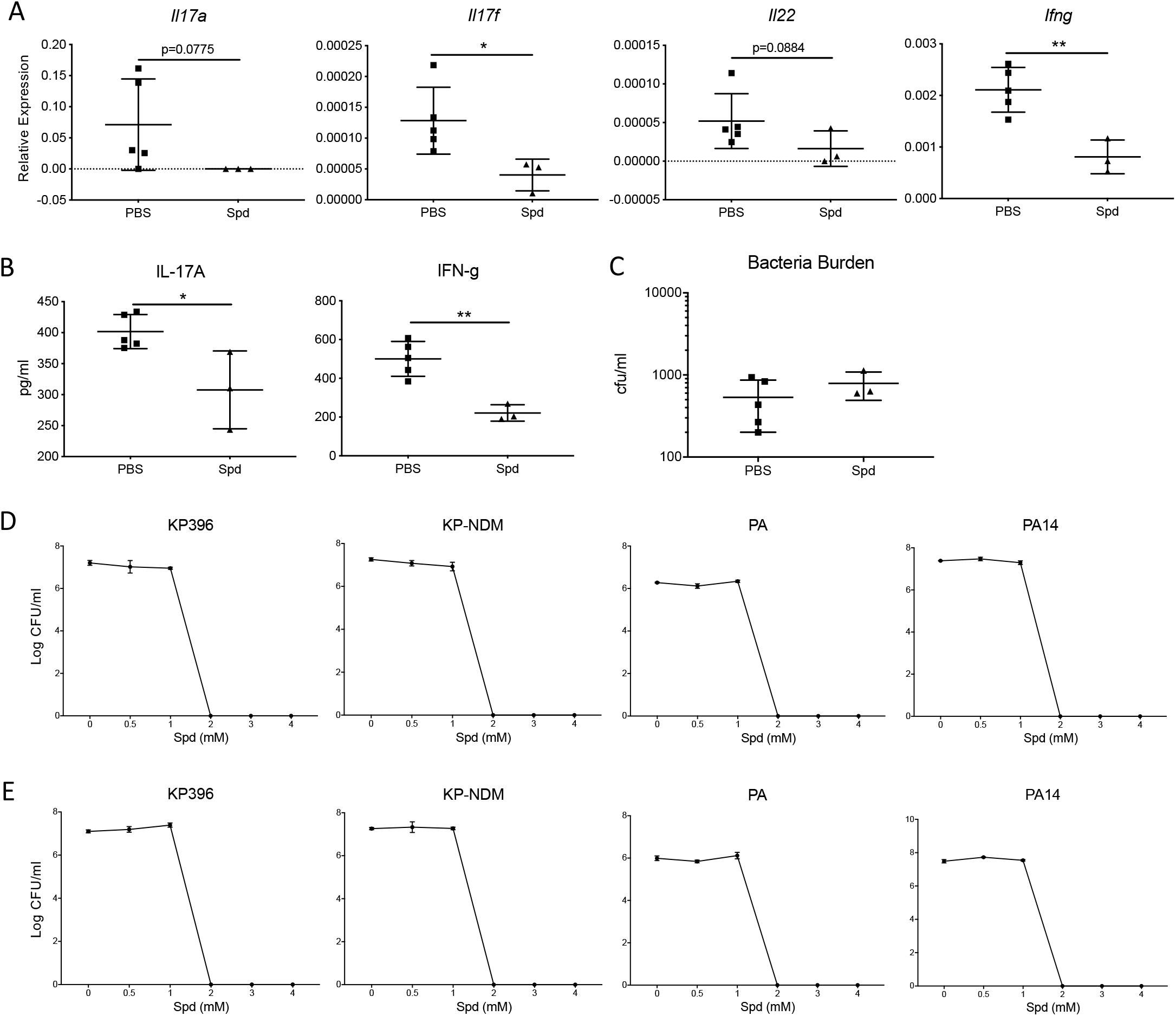
Spermidine treatment suppresses Th17/IL-17 pathway in a live PA bacteria infected mouse model. A: Spermidine alleviates Th17/IL-17 pathway production in PA infected mice model (N=5 in PBS group; N=3 in Spd group). Th17/IL-17 pathway gene mRNA and protein in lung tissue were achieved by RT-PCR and ELISA, relative expression to Hprt. B: Lung PA bacteria burden (K) was obtained with cfu assay. This experiment has been repeated once. C: CFU assay to determine bactericidal activity of varying concentrations of spermidine after 4 hours. Live bacteria including KP396, KP-NDM, PA (as PA ATCC10145) and PA14 were treated with 0, 0.5, 1, 2, 3 and 4mM spermidine. D: CFU assay to determine bactericidal activity of varying concentrations of spermidine after 24 hours. Live bacteria including KP396, KP-NDM, PA (as PA ATCC10145) and PA14 were treated with 0, 0.5, 1, 2, 3 and 4mM spermidine.

Upregulation of Th17/IL-17 pathway expression is expected to enhance bacterial clearance, while inhibiting Th17/IL-17 signaling can lead to bacterial retention.^21,22^ Inhibiting the expression of Th17/IL-17 pathway with spermidine may therefore hinder bacterial clearance in mice, posing a potential concern for elevated infection risk during inflammation management. However, the quantitative measurement of residual bacteria burden in the lungs showed that spermidine did not adversely affect the clearance PA bacteria (Figure 4C). To further investigate this unimpaired bacterial clearance, we directly administered spermidine to different subtypes of KP (KP396, KP-NDM) and PA (PA ATCC10145, PA14) *in vitro*. Both KP and PA were completely eradicated after 4 hours of spermidine treatment at spermidine concentrations of ≥2mM (Figure 4D). The bactericidal effect of spermidine after 24h of treatment is the same as that of after 4h (Figure 4E). Direct bactericidal effect of spermidine may aid in bacterial clearance, maintaining mucosal barrier homeostasis despite the downregulation of Th17/IL-17 pathway expression.

To further determine whether spermidine targets T cells for reducing inflammation, we used T cell knockout mice to explore whether spermidine acts on other immune cells. We established LPS-induced acute lung injury and live bacterial infection models in Rag2/II2rg double gene knockout mice, which are deficient in T cells, B cells and NK cells.^24^ The Th17 cytokines were completely absent in these mouse according to RT-PCR (data not shown), confirming that T cell function were knocked out. According to both ALI (Figure 5A-C) and PA14 infection (Figure 5D-F) models, spermidine did not protect against inflammation in Rag2/II2rg double gene knockout mice. All markers of inflammation were elevated with or without spermidine administration, including the number of BAL immune cells, the proportion of BAL monocytes, the proportion of BAL neutrophils, BAL total protein, wet-to-dry specific gravity, edema, and lung histopathological score (Figure 5). This stark loss in spermidine’s anti-inflammatory effect accentuates that spermidine is likely acting upon Th17 T cells in the Th17/IL-17 pathway and not through other immune cells.

**Figure 5.**
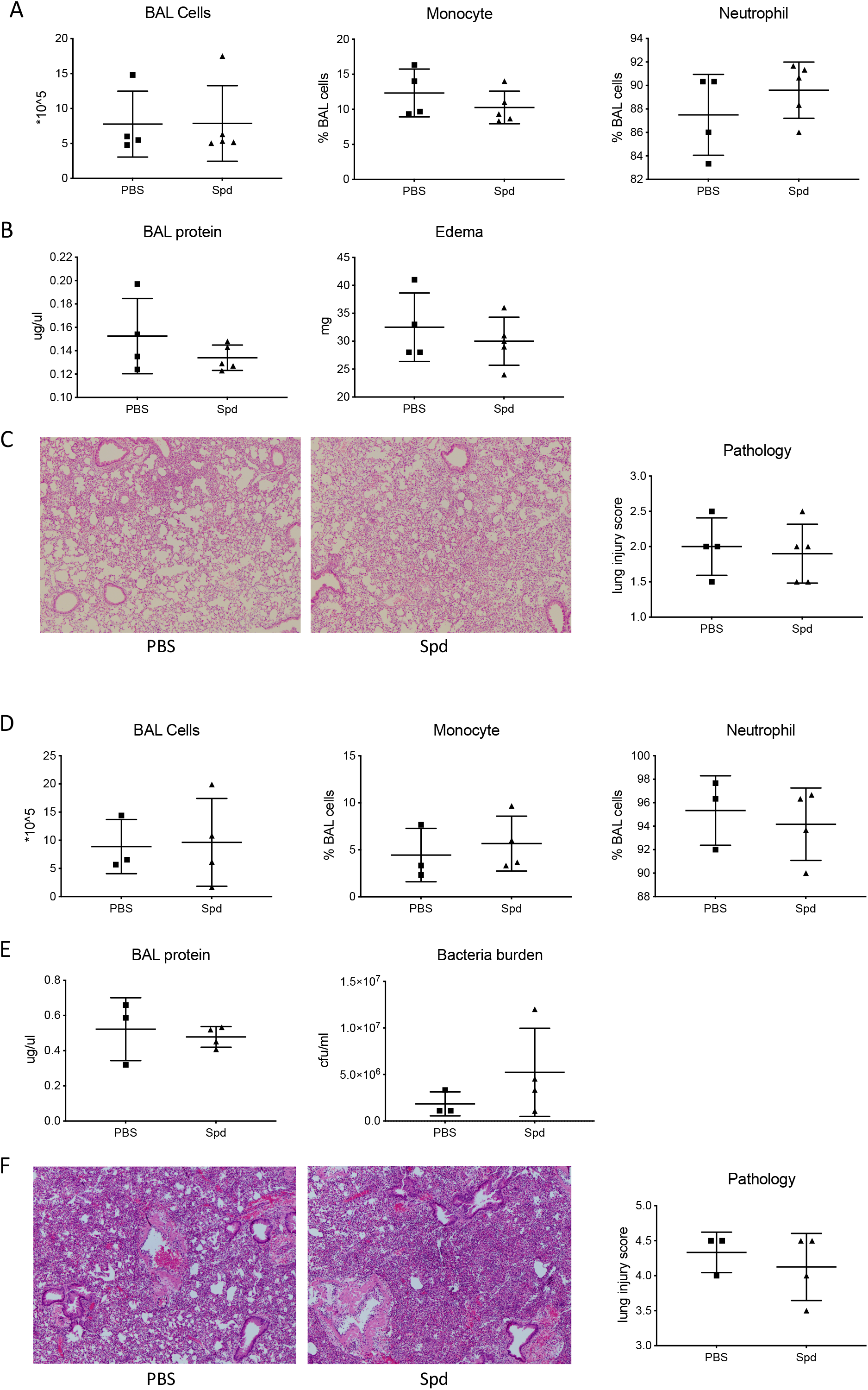
Lymphocyte depleted Rag2/Il2rg double KO mice do not respond to spermidine. 6-8 weeks Rag2/II2rg double knockout mice were injected with Spermidine 50 μg/g according to mouse weight or PBS by IP twice at −24 h and 0 h separately. Then mice from both groups were inoculated with 5 μg/g LPS (A-C) according to mouse weight or 1*10^6 cfu PA14 live bacteria (D-F) by IT at 0 h. 24 h later, BAL and lung tissue were harvested for downstream analysis. A-C: Immune cell differentiation (A), BAL protein and lung tissue edema (B) histology (C) were obtained in LPS induced ALI model; D-F: In live bacterial infection model, immune cell differentiation (D), BAL protein, bacteria burden (E) and pathology (F) were also assessed.

### Spermidine down-regulates Th17/IL-17 cytokine gene expression by regulating PRDX1 activity

To determine the intrinsic mechanism of how effector Th17 cytokines are downregulated by spermidine in T cells, we performed scRNA-Seq sequencing and bulk mRNA sequencing analysis on IL-1β/IL-23 activated splenic T cells (SPT) and lymph node (LN) cells treated with spermidine. To minimize batch effect between samples, we multiplex all the samples with cell hashing method^23^ for spermidine treatment (SPT sper and LN sper) and untreated groups (SPT stim and LN stim) (Figure 6A). The results after de-multiplexing^24^ show that, compared with the spermidine untreated T cell control group (SPT stim), the expression of *Txn1* and *Prdx1* genes were significantly increased in the spermidine treatment group (Figure 6B-C). *Prdx1* encodes peroxidase 1 (PRDX1), which is an antioxidant enzyme that protects the cell by reducing hydrogen peroxide retention.^25, 26^ Coincidentally, thioredoxin (TXN1) is an oxidoreductase also with antioxidant activities^27, 28^. Other genes such as *Txnrd1* and *Stat1* did not change significantly. We further confirmed with mRNA-Seq data that the expression of *Txn1* and *Prdx1* genes in T cells in the spermidine treatment group was increased directly in proportion to the concentration of spermidine (Figure 5D). In the subsequent RT-PCR verification, we observed that only the *Prdx1* gene was highly expressed in the spermidine treatment group (Figure 5E). Prdx1 activity has an especially strong association with spermidine treatment, raising the possibility of Prdx1 being a key spermidine effector.

**Figure 6.**
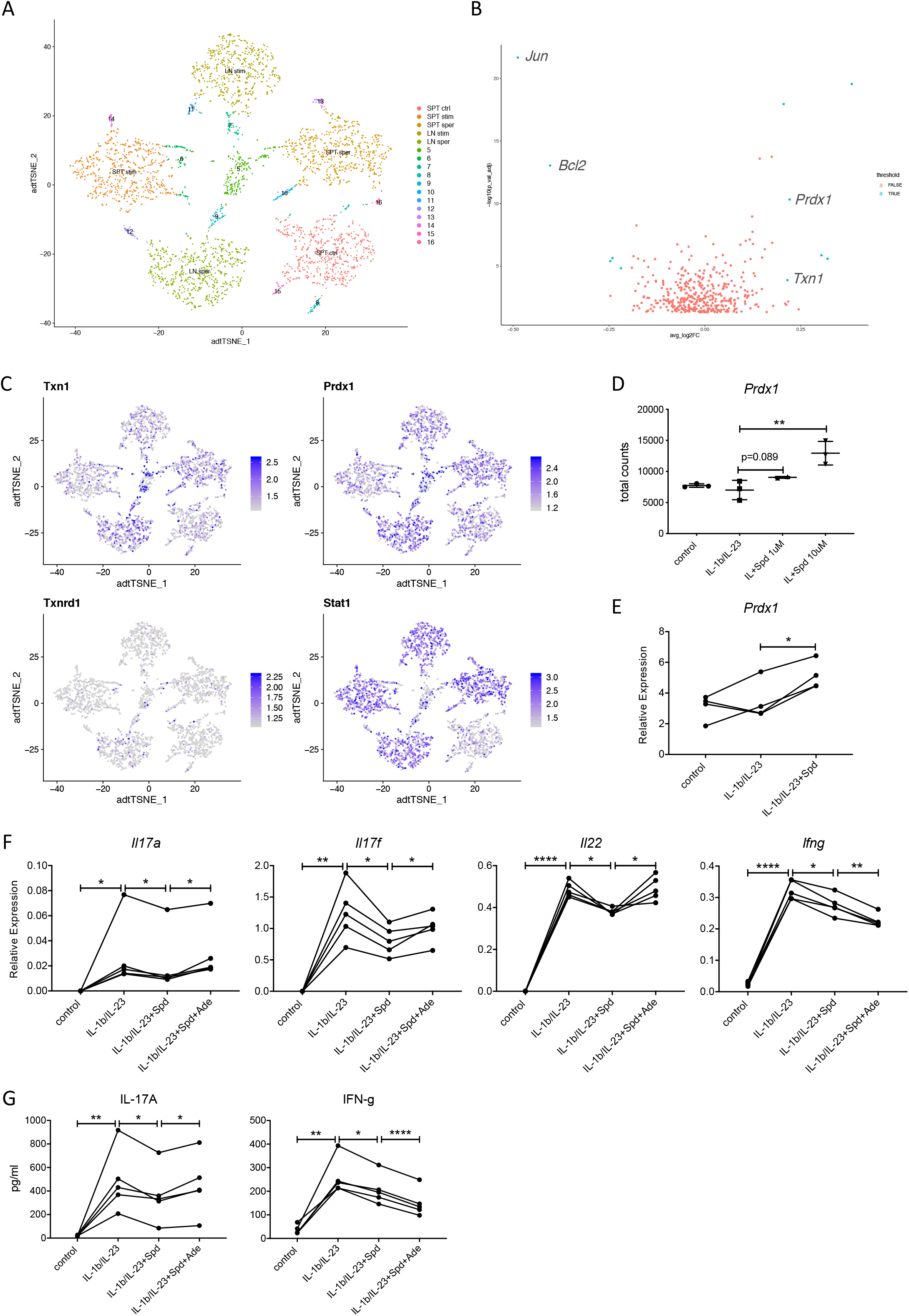
Spermidine inhibits IL-17 production in a *Prdx1* dependent manner. A: scRNA-Seq integration and cell cluster label of fresh lymph node cells and enriched spleen T cells, which were stimulated with mouse IL-1b +IL-23 50ng/ml at the presence or absence of 3uM Spermidine treatment overnight. Cells were collected for scRNA-Seq performance. 5 major cell population was identified by scATAC-seq data: spleen T cell control group (SPT ctrl); spleen T cell IL-1b/IL-23 stimulation group (SPT stim); spleen T cell IL-1b/IL-23 plus spermidine treatment group (SPT sper); lymph node cell IL-1b/IL-23 stimulation group (LN stim); lymph node cell IL-1b/IL-23 plus spermidine treatment group (LN sper). B: Antioxidant activity related genes Txn1 and Prdx1 were found significantly upregulated after spermidine treatment to IL-1b/IL23 activated IL-17 producing cells. C: Volcano plots representing differentially expressed genes between IL-1b/IL-23 stimulation group vs IL-1b/IL-23 plus spermidine treatment group (cutoffs: LogFC=0.2 and padj=0.001). D-E: Both mRNA-seq data (D) and RT-PCR (E) revealed *Prdx1* as a potential target for spermidine, relative to *Hprt* expression, N=3. F-G: Spermidine inhibits IL-17 production in a *Prdx1* dependent manner. Fresh enriched spleen T cells were stimulated with 50ng/ml mouse IL-1b +IL-23, 50ng/ml IL-1b/IL-23 + 3uM Spermidine, 50ng/ml IL-1b/IL-23 + 3uM Spermidine + 10uM Prdx1 inhibitor Adenanthin (Ade) separately, incubating overnight. Ade treatment reversed the suppression of Th17/IL-17 pathway induced by spermidine. *Il17a*, *Il17f*, *Il22* and *Ifng* gene expression was measured by qPCR (F), relative expression to *Hprt*; IL-17A and IFN-g proteins were tested by ELISA (G). This figure is a representative of four experiments, N=2-3 in each experiment.

To determine whether spermidine regulates Th17 cytokine production through PRDX1, adenanthin was co-administered with spermidine into activated Th17 cells. Adenanthin is a recently discovered inhibitor against antioxidant enzymes, including PRDX1, derived from a plant extract^29, 30^. As previously observed, IL-1β/IL-23 activated the Th17/IL-17 and IFN-γ pathways, while spermidine significantly down-regulated Th17/IL-17 and IFN-γ products (Figure 5F). Compared to spermidine treatment alone, the addition of adenanthin enhanced the gene expression of *Il17a*, *Il17f*, *Il22* as well as the levels of Th17/IL-17 products (Figure 5F-G). Adenanthin successfully reversed the inhibitory effect of spermidine on Th17/IL-17 pathway, suggesting that spermidine down-regulates Th17/IL-17 pathway by regulating PRDX1 activity. Interestingly, the combination of adenanthin and spermidine treatment further inhibited the gene and protein expression of IFN-γ (Figure 5F-G), which needs to be further explored in future investigations.

Adenanthin and DFMO are both inhibitors against spermidine’s action against Th17/IL-17 signaling. DFMO inhibits upstream of spermidine action during spermidine synthesis, while adenanthin inhibits downstream of spermidine by inhibiting PDRX1. Administration of DFMO reverses the inhibitory effect of spermidine on the Th17/IL-17 pathway and enhances the expression of Th17/IL-17 pathway products. In contrast, coadminstration of adenanthin with spermidine did not completely restore the expression of Th17/IL-17 pathway products. This suggests that the spermidine’s activity is not completely inhibited by suppressing PDRX1 activity. These results indicate that spermidine may inhibit the expression of Th17/IL-17 products through additional mechanisms, which need to be investigated further.

## 2.4 Discussion

The key position of Th17/IL-17 signaling pathway in the body’s immune system highlights the importance of studying the regulatory mechanism of this pathway. Understanding the regulation mechanisms of the Th17/IL-17 signaling pathway, especially the regulatory factors of Th17 cells, have always been of great interest for immunology research. Th17 cells have been reported to be inhibited by cytokines secreted by most other types of T cells such as Th1 cells and Th2 cells, ^31, 32, 33^ which have important roles in controlling inflammatory cascades and hyper-immunity. Regulatory T cells play a role in monitoring and regulating Th17 cells, but the specific regulatory mechanisms for the Th17/IL-17 signaling pathway has been elusive.^34, 35^

Polyamines, especially spermidine and spermine, have been implicated with inflammation, with elevated levels of polyamines found in the inflammatory sites for infection, trauma, tumor and autoimmune diseases.^36, 37^ Prior to this study, the mechanism through which polyamines enact their anti-inflammatory effects had not yet been established. In this study, we established a key regulatory mechanism of spermidine on Th17/IL-17 signaling pathway. Based on both *in vitro* and *in vivo* models, we found that spermidine directly acts on T cells to down-regulate the Th17/IL-17 signaling pathway. Based on RNA sequencing data, we determined that spermidine down-regulates the Th17/IL-17 signaling pathway by controlling PRDX1 protein activity in T cells. PRDX1 protein is mainly involved in antioxidant functions and helps to remove reactive oxygen species (ROS) in cells. PRDX1 activity is important during times of cellular oxidative stress, such as in inflammation, where excessive amounts of reactive oxygen species are produced in the cell.^38,39^ Our observation that spermidine acts through PRDX1 is consistent with the literature that spermidine is involved in controlling oxidative stress.^40,41,42^

We also found that spermidine treatment, despite its immune suppressive role, does not negatively affect lung bacterial clearance in mice. In addition to its anti-inflammatory and immunomodulatory effects, spermidine was found to have a direct bactericidal effect against PA and KP. These results suggest that spermidine may be key factor in the host-infection interface, maintaining mucosal barrier homeostasis by controlling infection without detrimental levels of inflammation. Further understanding of how spermidine endogenously controls the dynamic balance between Th17/IL-17 suppression and infection will guide clinical strategies against infection induced hyper-immunity.

Further investigations that explore other molecular mechanisms by which spermidine inhibits the Th17/IL-17 are necessary. Our comparison of DFMO and adenanthin, inhibitors of spermidine synthesis and effectors, respectively, indicate that spermidine may have additional Th17/IL-17 pathway effectors besides PRDX1. In addition, whether other polyamines, which also have been implicated with anti-inflammatory effects, ^36, 37^ engage in a similar regulatory mechanism as spermidine must also be investigated. Furthering the mechanistic understanding of Th17/IL-17 regulation by endogenous and exogenous spermidine may aid efforts to clinically manage debilitating cases of infection and autoimmune diseases.

## Materials and methods

### 1. Mouse models

All the mice used in this project were of wildtype strains and purchased from Jackson Lab (Cat# 000664). Animals were maintained in pathogen-free conditions at the core animal facility at the University of Pittsburgh Medical Center. Mice age matched to 6-8 weeks old were used to set up pre-designed models with the approval from the University of Pittsburgh Institutional Animal Care and Use Committee.

#### 1.1 Mouse peritoneal neutrophil induction and treatment

4% sterile Thioglycollate (Sigma, Cat# 70157) in ddH_2_O was applied to induce mice peritonitis, 100 μl per mouse with intraperitoneal (IP) injection. 5 h later, mice were sacrificed by CO_2_ euthanasia. Peritoneal lavage samples were collected by injecting sterile PBS into peritoneal cavity with a 10 ml syringe. Peritoneal lavage cells were isolated and stimulated overnight at 37 °C with 50 ng/ml m-IL-1β (Biolegend, Cat# 575002) plus m-IL-23 (Biolegend, Cat# 589002) in 10% FBS-IMDM media. Cells were collected for scRNA-seq analysis.

#### 1.2 Mouse lung inflammation induction and treatment

Either LPS (Sigma Inc.) or P.A. (ATCC® 10145GFP™) were inoculated into mice to mediate acute mouse lung inflammation. Spermidine (Sigma, Cat# S2626-1G) or the spermidine synthesis inhibitor DL-α-Difluoromethylornithine (DFMO, Cayman Chemical, Cat# 16889) were administered at time of inoculation to explore the roles of spermidine in inflammation regulation. Three modules were designed in this project: LPS administration (5 μg/g body weight) + spermidine treatment (5 μg/g body weight); P.A. infection (1×10^6^ cfu/mouse) + spermidine treatment (5 μg/g body weight); LPS administration (5 μg/g body weight) + DFMO treatment (1.5 mg/g body weight). For spermidine treatment modules, mice were injected intraperitoneally with spermidine solution or PBS as control, then anesthetized with isoflurane and challenged with LPS or PA intratracheally (IT). For DFMO treatment experiments, mice were pretreated with DFMO or PBS intraperitoneally for 1 hr, after which all mice were anesthetized with isoflurane and intratracheally inoculated with LPS. 18h after inoculation, mice from all the modules were euthanized with CO_2_ for further processing.

### 2. BAL and lung tissue processing

Before harvesting the lungs, mouse lungs were lavaged with sterile PBS through the trachea. BAL cells were obtained by centrifuging the BAL fluid at 300xg for 10 min at 4 °C. Right after centrifuging from the cytocentrifuge (Cytospin 4, Thermo Scientific), fresh BAL cell cytospin slides were stained using a Kwik-Dif Stains kit (Fisher Scientific, #99-907-00) for inflammatory cell differential counts. Right superior and inferior lobes were homogenized in 1 ml sterile PBS using gentleMACS Octo Dissociator (Miltenyi Biotec). A 100 μl aliquot of homogenate was reserved for bacteria cfu assay. The rest of the homogenate was centrifuged at 4 °C over 10000 g for 10 min, and the supernatant was stored at −80 °C for further use in ELISA, LDH assays, and other experiments. The right middle lobe was cut down and incubated in a 65 °C incubator for 48 h, and the wet/dry weight ratio as well as the edema levels were measured. The left inferior lobe was immersed in RLT buffer with 2-mercaptoethanol for RNA extraction. The left superior lobe was fixed with 10% neutral buffered formalin (Sigma, Cat# HT501128-4L) at 4 °C for over 24 h and subsequently subjected to H&E staining.

### 3. LDH detection

LDH Cytotoxicity Assay (Promega, Cat# G1780) was used to quantify LDH levels as a function of cytotoxicity. The procedure was strictly performed according to the manufacturer’s protocol.

### 4. Bactericidal experiments by spermidine

Live bacteria including KP396, KP-NDM, PA and PA14 were treated with 0, 0.5, 1, 2, 3 and 4mM spermidine for 4 h and 24 h respectively. After the given time points, bactericidal activity was determined by using a CFU assay.

### 5. Bacteria CFU assay

This assay was used to accurately assess bacteria burden. Briefly, the primary sample was serially diluted as 1:10, 1:100, 1:1000, 1:10000, 1:100000, 1:1000000. A10 μl aspirate from each diluted sample were transferred to a LB agar plate in technical triplicates. The agar plates were incubated at 37 °C overnight and the resulting bacteria colonies were counted carefully. The final CFU calculation formula was as followed: the bacterial concentration in the sample tested (CFU/ml) = (Total number of colonies from the triplicate dots /3) *dilution factor*100.

### 6. Primary T cell isolation

Fresh T cells from spleen and lung draining lymph node were used for in vitro experiments. Spleen and lung draining lymph nodes were harvested from WT mice after CO_2_ euthanasia. Single cell suspensions from spleen and lymph node tissue were collected separately by crushing the organs with sterile, chilled PBS through a 70 μm filter (Fisher Scientific, Cat# 22-363-548) and a syringe.

A mouse T cell isolation kit (STEMCELL, Cat# 19851) was used to enrich T cells from harvested spleen cells. Unwanted cells were negatively purified from spleen cell suspensions by incubating spleen cells with biotinylated antibodies directed against non-T cells. The unwanted cells were magnetically isolated from total spleen cells with a streptavidin-coated magnetic bead, enriching the spleen cell suspension with T cells.

### 7. Primary cell stimulation

Enriched spleen T cells and lung draining lymph node cells were plated (3-4×10^6^ cells) in a 24-well plate, using 0.5 ml IMEM, 10% FBS, and standard Penicillin-Streptomycin antibiotics in each well. Cells were treated with various interventional conditions, as followed: 50 ng/ml m-IL-1β/m-IL-23; 50 ng/ml m-IL-1β/m-IL-23+3 μM spermidine; 50 ng/ml m-IL-1β/m-IL-23+20 mM DFMO; 50 ng/ml m-IL-1β/m-IL-23+3 μM spermidine+10 μM Ade (ChemFaces, Cat# CFN99215). For DFMO experiments, DFMO and control PBS were added to the cells 4 h prior to IL-1β/IL-23 stimulation to block spermidine synthesis. For other experimental groups, all treatments were administered to the cells at the time of stimulation. At 18 h time point, cells were harvested for RNA-seq, spermidine quantification by HPLC, or lysed in RLT buffer with 2-mercaptoethanol for gene expression. The supernatant was used for ELISA experiments.

### 8. RNA Extraction and cDNA Synthesis

Cell and lung tissue RNA was extracted with RNeasy Miniprep Kit (QIAGEN, Cat# 74136), in accordance with the manufacturer’s instruction. cDNA was synthesized using qScriptTM cDNA Synthesis Kits (Quantabio, Cat# 95047-100).

### 9. Real-Time PCR

Real-time PCR was constructed and read by the Bio-Rad CFX96 system using TaqMan PCR Master Mix (Bio-Rad, Cat# 1725284) and pre-mixed probe sets: *Il17a* (Mm00439618_m1), *Il17f* (Mm00521423_m1), *Il22* (Mm01226722_g1), *Ifng* (Mm01168134_m1), *Srm* (Mm00726089_s1), *Smox* (Mm01275475_m1), *Prdx1* (Mm01621996_s1), *Gapdh* (Mm99999915_g1) and *Hprt* (Mm03024075_m1), from Thermo Scientific.

### 10. ELISA

ELISA kits were used to measure mouse IL-17A (BioLegend, Cat# 432506) and mouse IFN-γ (BioLegend, Cat# 430804). The procedures were strictly employed according to the manufacturer’s instructions.

### 11. ScRNA-seq, RNA-seq and Total-seq

ScRNA-seq libraries were constructed and sequenced following the “Single Cell 3’ Reagent Kits v2 User Guide” (10X Genomics). RNA-seq methodology was developed in close reference to a published study.^43^ Similar to ScRNA-seq, Total-seq was performed following the cell hashing protocol (BioLegend). Briefly, five groups of cells were pre-labelled with mouse HashTag oligos (HTO) on cell surface: spleen T cell control group (SPT ctrl) with hashtag 1 (BioLegend, Cat# 155801); spleen T cell IL-1β/IL-23 stimulation group (SPT stim) with hashtag 2 (BioLegend, Cat# 155803); spleen T cell IL-1β/IL-23 plus spermidine treatment group (SPT sper) with hashtag 3 (BioLegend, Cat# 155805); lymph node cell IL-1β/IL-23 stimulation group (LN stim) with hashtag 4 (BioLegend, Cat# 155807); lymph node cell IL-1β/IL-23 plus spermidine treatment group (LN sper) with hashtag 5 (BioLegend, Cat# 155809). The five cell populations were mixed and barcoded, and barcoded cDNA were synthesized via reverse transcription. Further cDNA libraries were constructed by cDNA amplification. HTO-derived cDNAs (<180bp) and mRNA-derived cDNAs (>300bp) were separated using a magnetic beads selection methodology. An HTO sequencing library was generated for sequencing with another round of cDNA amplification. mRNA-derived cDNA library was constructed following the conventional scRNA-seq protocol. HTO library and DNA library were then sequenced on an Illumina HiSeq sequencer (MedGenome, Inc.). Sequencing data was initially analyzed by Cell Ranger (10X Genomics) and downstream analysis was done by R.

### 12. Statistics

All data analyses were achieved and graphed by Prism 8.0 (GraphPad). The one-way ANOVA test was performed to compare differential gene expression among four groups. Paired Student’s t-test was applied for other comparisons between the paired two groups,

## Data Availability

R code used for RNA-seq analysis is available upon request. The sequencing data reported in this paper will be deposited to the Gene Expression Omnibus.

